# Genome report: Genome sequence of the hibiscus mealybug, *Nipaecoccus viridis* (Newstead), an invasive pest of citrus

**DOI:** 10.1101/2025.02.06.636937

**Authors:** Tracy Liesenfelt, Amanda Markee, Emilie P. Demard, Lauren M. Diepenbrock, Andrew J. Mongue

## Abstract

Mealybugs are frequently known for being pest insects to both ornamental and large-scale agricultural crops. Yet despite their agricultural importance, the genomic resources for this group remain quite small. One such species is the hibiscus mealybug, *Nipaecoccus viridis* (Newstead) (Hemiptera: Coccomorpha: Pseudococcidae). This invasive mealybug species has recently expanded throughout Florida and has spread across the state. Genomic resources would provide a new means to better understand the invasive nature of this insect, and thus, we present the de novo genome assembly for *Nipaecoccus viridis*. Our genome assembly is 289 Mb, in which 91.2% of this sequence assembled into 5 chromosomal scaffolds. We report 15,370 genes to be present within our genome. We found that repetitive elements in the genome accounted for 32.40% of the sequence. These statistics follow similar trends to other previously sequenced pseudococcid species.

## Introduction

While often small and unnoticed, mealybugs (Hemiptera: Coccomorpha: Pseudococcidae), can pose large threats to agricultural systems and inflict economically important damage to crops. Paired with many species’ highly invasive nature, this group of insects is important to study, not just for its agricultural importance, but from the basic research perspective as well (Liebhold et al. 2024). Scale insects which include the mealybugs, exhibit many unique modes of sex determination, including the poorly understood system of paternal genome elimination (PGE), a rare form of haplodiploidy (Nur 1966; de la Filia et al. 2015). Under PGE, while males will inherit their genetic material from both parents, they are only able to pass on their mother’s genes to their offspring (Herbette and Ross 2023). Within mealybugs, one set of the genome is deactivated. This occurs through heterochromatinization within their early development, and while this inactive set can still be divided into somatic cell lines, it is not passed onto their offspring as it is destroyed in meiosis before it can end up in mature sperm (Nur 1990; Mongue et al. 2024). Using the present technologies, the assembly of genomes within the family Pseudococcidae holds the key to understanding these insects at a much deeper level than previous studies.

The hibiscus mealybug, *Nipaecoccus viridis* (Newstead), is one such insect within this family. As adults, these insects are highly sexually dimorphic (**Figure 1)** having winged males with a body length of around 1.3 to 2.5 mm and adult females have soft oval-shaped body about 2.5 to 3 mm and 1.5 to 2.5 mm wide (Sharaf and Meyerdirk 1987). Adult females use waxy secretions to create an ovisac beneath their bodies to harbor their eggs; a single female produces anywhere between 400 to 1,000 eggs within her lifetime (Olabiyi, Middleton, et al. 2023).

**Fig. 1.**
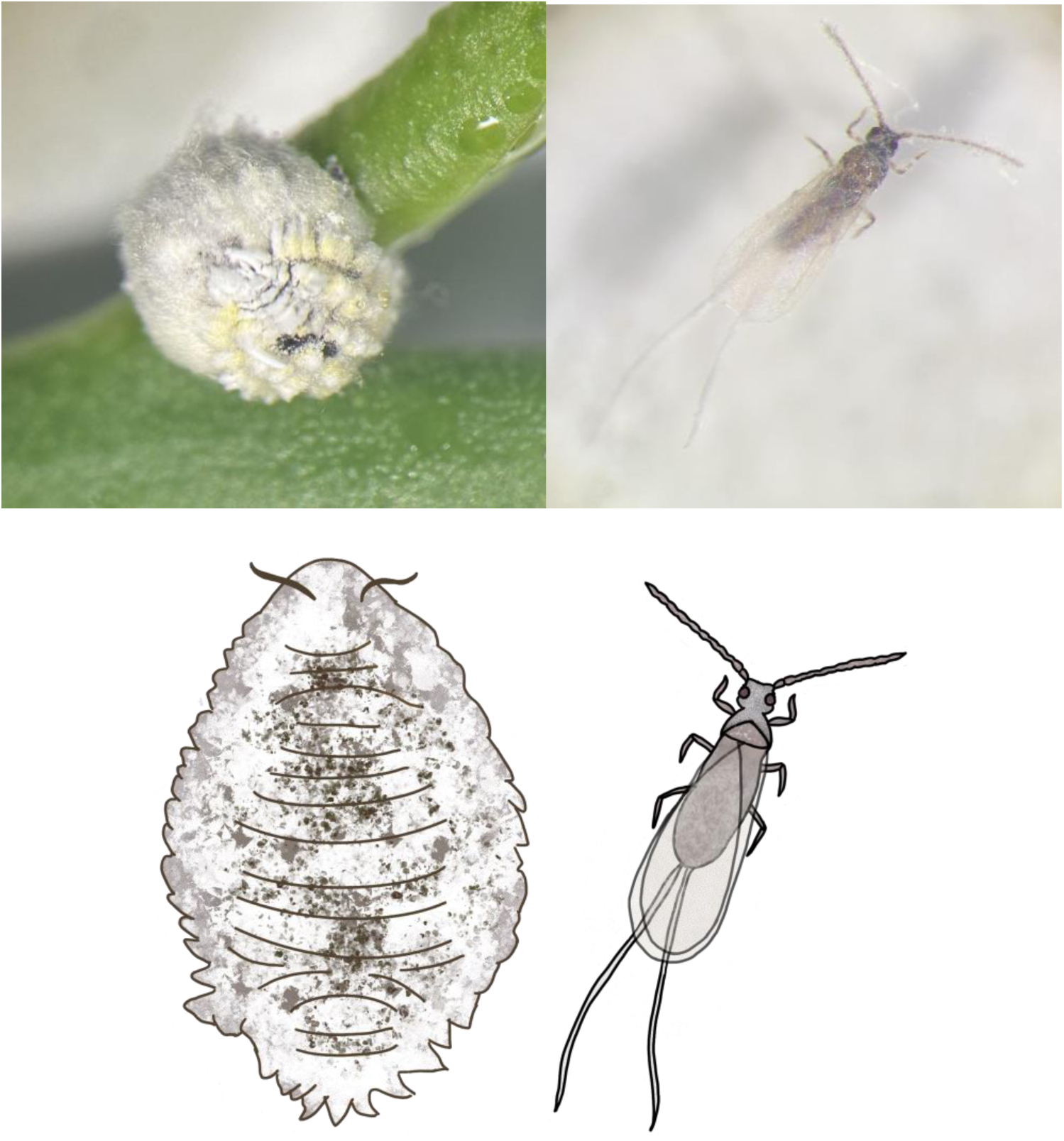
Depictions of sexual dimorphism present in adult *Nipaecoccus viridis*. **Top Left**: An adult female covered in wax with her ovisac underneath her body. **Top Right**: A winged adult male. **Bottom**: Illustrations depicting the adults of *Nipaecoccus viridis* to highlight their sex differences. Photos by TL, illustrations by TL.

While their ancestral geographic range is not known, these insects are believed to have originated in Asia (Evans and Dooley 2013) and have since been spread to at least 60 different countries (Garcia et al. 2016). Part of this worldwide invasive success comes from their massive host range. *Nipaecoccus viridis* is highly polyphagous, feeding on at least 140 recorded genera of plants (Garcia et al. 2016). Within this expansive list, this insect can be a major pest on several economically important crops including soybeans, pomegranate, cotton, cassava, and citrus, the latter being important to the recent invasion of *N. viridis* into Florida.

In 2009, *N. viridis* was recorded in a natural area in Palm Beach County, Florida (Stocks and Hodges 2010). It was not until 2019 when a second instance of the mealybug was recorded in commercial citrus found in Highlands County that concerns were raised regarding the damage it could pose to Florida’s agriculture (Diepenbrock and Ahmed 2020). These insects pose a threat to not just the plants, but the marketability of the fruit produced. In particular, *N. viridis*, can cause a range of damage when feeding on citrus, including fruit drops, deformed fruit (**Figure 2**), and even death to young trees (Diepenbrock and Ahmed 2020; Olabiyi et al. 2023).

**Fig. 2.**
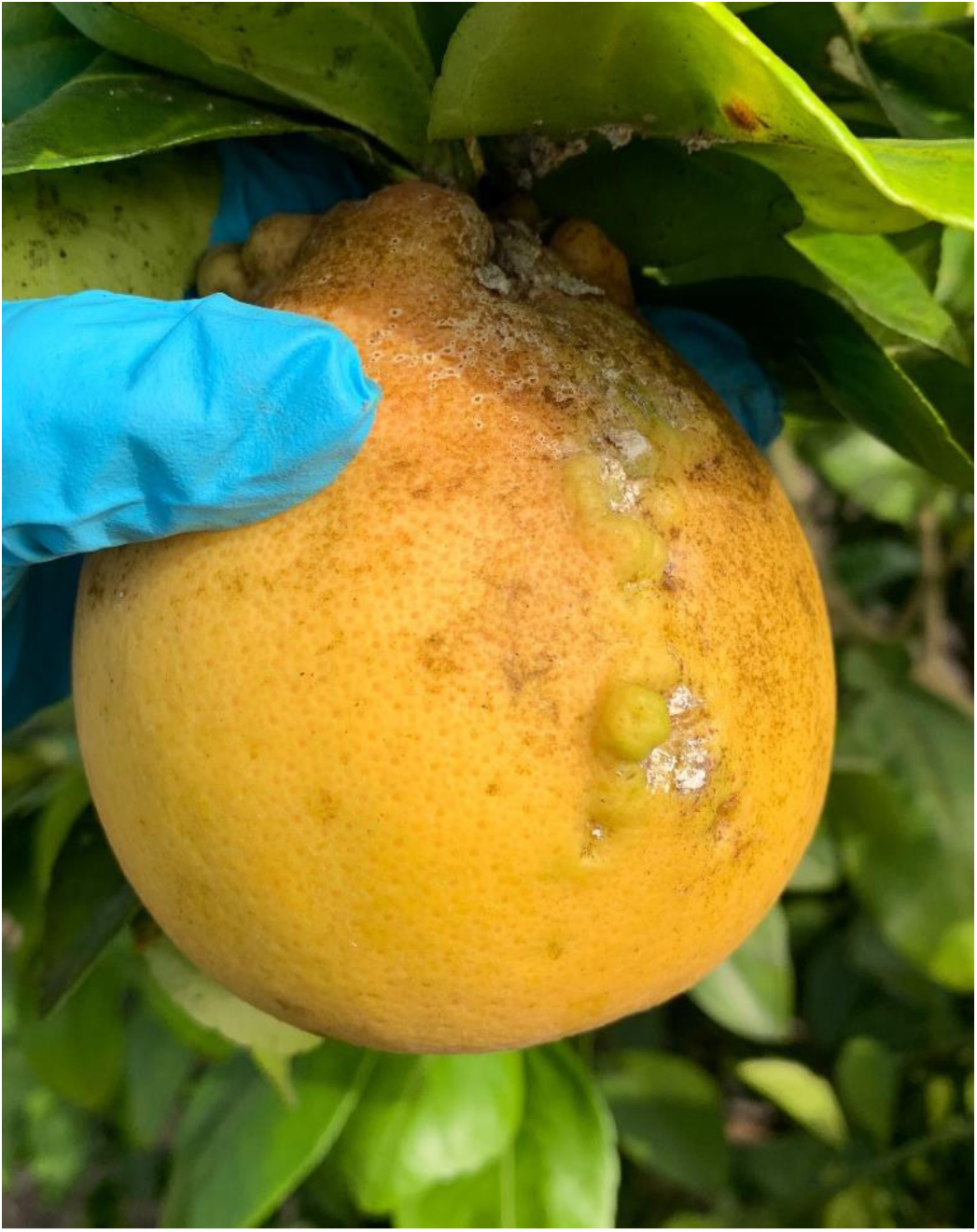
Pictures of deformations present on a grapefruit due to feeding damage from *Nipaecoccus viridis.* Their feeding causes gall-like outgrowths on the surface of the fruit. Along with the outgrowths, residual white wax can be seen left behind on the fruit. Photo by TL.

The origins of this invasion into Florida are unknown, but since the first detection in commercial citrus in 2019, *N. viridis* has become widespread across the southern region of the state, now having been recorded in 24 different counties (Deeter and Ahmed 2023; Olabiyi et al. 2023; and personal communication with Florida Department of Agriculture and Consumer Services). Observation data alone cannot resolve invasion history, but genomic tools like a reference genome will allow researchers to explore the population history through variations in genetic markers. To facilitate this ultimate goal, as well as provide more resources on a rare sex determination system, this study presents the first chromosome-level assembly of *N. viridis*.

## Materials and Methods

### Sample Collection

DNA was extracted from adult female *Nipaecoccus viridis* collected from a lab-reared colony. This colony was initially established in 2019 from a field-collected population taken from Highlands County, FL (latitude: 27°20’24.00” N; longitude:-81°20’24.00” W) and reared in an indoor insectary at the University of Florida Citrus Research and Education Center (CREC) in Lake Alfred, FL. This colony was maintained at the temperature of 25 ± 5 °C, a relative humidity of 70 ± 10%, and a photoperiod of 16:8 h [L:D] following the conditions established in Olabiyi et al. (2023). The colony has been maintained without additional introductions from the wild population and has since become a distinct-inbred line. For our genome project, adult females were collected in August 2023 from this colony, roughly 45 generations after the initial establishment based on the approximately 32-day life cycle. Under the mealybug sex determination system of paternal genome elimination, both males and females share the same set of chromosomes across the genome, i.e. there are no sex chromosomes, (Nur 1990) therefore, we chose females for the convenience of larger body size than males with no tradeoff in chromosomal sampling. Whole live females were flash frozen in liquid nitrogen and then stored in a-80 °C freezer prior to extraction.

### DNA Extraction and sequencing

To obtain DNA for sequencing, we homogenized individual females using a PowerMasher II tissue disruptor and then we extracted the DNA using an Omniprep DNA extraction kit (G-Biosciences). We used the standard Omniprep base kit protocol, with minor adjustments following protocols from Mongue et al. (2024), which successfully generated a chromosome-level assembly of another scale insect species. Specifically, tissue was lysed overnight, but no longer than 16 hours on an incubator at 56°C. During the DNA precipitation step, 2 microliters Mussel Glycogen were added to each sample to increase DNA yield. Once this reagent was added to each sample, we allowed DNA to precipitate for 1 hour in a-20°C freezer. These modifications allowed us to maximize our DNA yields and decrease contamination of samples due to the wax coating on the individual females being extracted. We checked our extractions using a combination of Qubit Fluorometer with Qubit dsDNA broad range Sensitivity Assay Kit and Nanodrop spectrophotometer for DNA concentration and quality, respectively.

After quality control of our genomic DNA, we submitted a pooled extraction of 9 adult females from the inbred colony to be sequenced at the University of Florida Interdisciplinary Center for Biotechnology Research (ICBR) on a PacBio Sequel IIe platform (Menlo Park, Ca, USA) for HiFi long read sequencing. We separately pooled 500 mg of whole adult female body tissue, collected from the same lab-reared colony to flash freeze in liquid nitrogen and submitted to Novogene Inc (Sacramento, CA, USA) for the generation of Illumina Hi-C linked reads using Illumina NovaSeq technology. In both cases, pooling across multiple individuals was necessary to meet quantity thresholds for library preparation but, thanks to the inbred lab colony source, did not introduce genetic variation from unrelated individuals.

### Genome size estimates

After pooling for sequencing, we did not have enough tissue for flow cytometry methods to estimate genome size. Instead, we used Jellyfish v. 2.2.4 (Marçais and Kingsford 2011), a tool that allowed us to estimate our expected genome size using the raw reads from PacBio by generating our k-mer frequencies. Data was analyzed using custom scripts in R studio v. 4.2.0 (R Core Team 2018).

### Genome Assembly

Our primary genome was assembled on the University of Florida HiPerGator cluster using HiFiasm v.0.18 (Cheng et al. 2021) with the-l 3 parameter to aggressively remove haplotigs. After this initial assembly, we used BlobTools v.1.0 (Laetsch and Blaxter 2017) to search for potential contaminants and endosymbionts, as seen in other mealybug species. Mealybugs typically contain a specialized organ known as a bacteriome where they house endosymbiotic bacteria that aid them in obtaining their essential nutrients that they cannot synthesize for themselves (Husnik et al. 2013; Garber et al. 2024), so we anticipated the presence of bacteria contigs in this assembly. These endosymbiont candidates were compared against the NCBI BLAST database to provide more specific identification of bacterial species but not removed at this stage of the assembly. At this stage, we also analyzed the assembly for genome completeness using analysis of the hemipteran ortholog dataset (hemiptera_odb10) of BUSCO v. 5.7.0 (Manni et al. 2021).

To achieve a chromosome-level assembly, we used the results of our Hi-C sequencing received from Novogene in combination with the PacBio primary assembly. We aligned the Hi-C reads to the primary assembly using the Arima Hi-C pipeline (https://github.com/ArimaGenomics/mapping_pipeline) and used Picard (https://broadinstitute.github.io/picard/) v.2.18.2 to check for and remove duplicates in the assembly. The purged assembly was then scaffolded using YaHS v.1.1 (Zhou et al. 2023). We generated an initial scaffolded assembly and created a Hi-C contact map through the generation of a JBAT file for visualization in Juicebox v.2.3.0 (Robinson et al. 2018). This assembly was manually curated and corrected for misalignments and updated with YaHs to produce a final assembly fasta. We again used BlobTools to identify bacterial sequences and BUSCO to assess the completeness of our curated assembly.

### Gene Annotation

For our assembly, we used a *de novo* gene annotation approach with Helixer (Holst et al. 2023). This is a novel machine-learning tool which only required our genome sequence and a general lineage as input; repeat masking and RNA evidence are not required for this tool. For our run, we used ‘invertebrate’ as the lineage. To assess this annotation, we ran BUSCO (in protein mode) using the hemiptera_odb10 dataset.

### Repeat Masking

For the characterization of repeats in our genome assembly, we used a combination of tools. We first use RepeatModeler v.2.0 (Flynn et al. 2020) to model de novo repeats by searching for long terminal repeats under the “-LTRStruct” parameter. This provided us with a generation of repeats unique to *N. viridis* which were then added to a custom library of repeats. This custom library included repeats from the 2020 Repbase arthropod and hemipteran repeat databases (Bao et al. 2015), and other repeats recorded in the genomes of different scale insects, also identified through Repeatmodeler. These additional insects include cottony cushion scale, *Icerya purchasi* (Maskell) (Mongue, Ross, et al. 2024), citrus mealybug, *Planococcus citri* (Risso 1813) (Ross et al. 2024), the Chinese wax scale, *Ericerus pela* (Chavannes) (Yang et al. 2019) and the tuliptree scale, *Toumeyella liriodendri* (Gmelin) (Mongue et al. 2024). This custom database was imported into RepeatMasker v.4.0.9 (Smit et al. 2019) for the generation of a soft-masked assembly and the summary of our repetitive elements.

### Identification of Endosymbionts

To identify potential co-bionts, other sequenced organisms, especially endosymbiotic bacteria, present within the genome assembly, we evaluated several pieces of evidence at two stages of assembly. First, we analyzed the primary assembly using BLAST+ v.2.9 with the tool Megablast (Chen et al. 2015) to quickly compare the assembly against known sequences in the NCBI BLAST database. Next, we aligned the raw PacBio reads to the primary assembly using Minimap2 v.2.28 (Li 2018) and sorted the alignment by coordinates with SAMtools v.1.9 (Danecek et al. 2021). We passed the resulting sorted bam file and the MegaBLAST output into the tool BlobTools v.1.0 (Laetsch and Blaxter 2017) to examine the contigs for taxonomic identification, coverage, and GC content. In particular, we were interested in contigs labeled as “Pseudomonadota,” showing higher coverage than the core genome, and distinct GC content. We then took the genomic sequences of these contigs and manually BLASTed them against the NCBI database to identify the specific bacteria species. Putative bacterial contigs were recorded as those that hit as bacteria via BlobTools annotation, and those that were assembled as circular DNA (denoted with a ‘c’ at the end of the contig number), or both (**Table 3**, columns 1 and 2).

Since some of these hits represent distinct bacterial genetic sequences, while others may be the result of horizontal gene transfer from bacteria to mealybug (Husnik et al. 2013), we did not remove any contigs at this time. After Hi-C scaffolding and manual curation steps previously listed, we repeated the BlobTools and manual BLAST process on the final curated scaffolded assembly. We tracked whether sequences flagged as bacterial were incorporated into the core chromosomal scaffolds (as expected of horizontally transferred genes) or remained as distinct sequences (as expected of true symbiont genomes). Finally, we used NCBI’s contaminant screening tool during assembly upload as a final piece of evidence for co-biont sequences. We removed the latter prior to archiving the genome sequence for public use but present those symbiont sequences here as a supplemental file.

## Results and Discussion

### Initial Genome Estimates

For our PacBio HiFi long reads, we received 2 runs of the same genomic library sequenced on 2 separate SMRTbells, the first yielding 4.5 Gb of read data and the second run yielding 2.2 Gb of read data for a combined total of 6.7 Gb of read data. We found 1 peak in our curve at a sequencing depth of ∼10x and estimated the genome to be 340 Mb using the following equation: Genome Size = the abundance of each k-mer (x-axis) * frequency (y-axis) / mean coverage (peak of the histogram). There is a steep decline until reaching a coverage depth of 4, which represents the erroneous k-mers under the curve, which are k-mers that contain errors from sequencing.

While some scale insects can have massive genomes, such as the cottony cushion scale, *Icerya purchasi*, with a genome over 1000 Mb (Mongue, Ross, et al. 2024), mealybug species typically range from 300 – 400 Mb. This can be seen in citrus mealybug, *Planococcus citri*, which has a genome size of 403.6 Mb (Ross, et al. 2024) and the New Zealand flax mealybug, *Balanococcus diminutus* (Leonardi), which has a size of 313.1 Mb (Ross, et al. 2024), and fall in the larger end of this range. Other species like the cotton mealybug, *Phenacoccus solenopsis* (Tinsley, 1898), have a more comparable size at 292.5 Mb (Li et al. 2020) to that of *N. viridis*, putting our genome size estimates for *N. viridis* to be in the lower range of the expected genome size for a mealybug species.

### Sequencing and Assembly

Chromosome number is consistent within Pseudococcidae, typically n = 5 (as reviewed in Gavrilov-Zimin et al. 2015) including *N. viridis* (Parida et al. 1982; Nur et al. 1987). As expected, based on this cytogenetic work, we assembled 5 haploid chromosomes. Following our final curation, we assigned 91.2% of the assembly to our 5 chromosomal scaffolds. In our final curation steps, we made 4 manual adjustments to our assembly using the Juicebox v.2.3.0. tool (Robinson et al. 2018).

### Gene Annotation

We identified a total of 15,370 protein-coding genes within our genome for *N. viridis* by using the Helixer tool (Holst et al. 2023). This annotation showed similar and in fact marginally better completeness to the BUSCO search of the genome assembly itself. Overall, we obtained 90.6% completeness (86.3% single copy, 4.3% duplicated), 1.5% fragmented, and 7.9% missing. By this metric, Helixer appears to have performed at least as well as more traditional evidence-based annotation methods used in other scale insects. By comparing *N. viridis* to other members of Pseudococcidae, we find that our recorded number of genes is on the lower end. Using the web resource Mealybug Base (https://ensembl.mealybug.org/index.html), we can note that *Planococcus citri* was recorded to have 40,620 genes and the longtailed mealybug, *Pseudococcus longispinus* (Targioni Tozzetti) was recorded to have 20,766 genes. Another scale insect, the tuliptree scale, *Toumeyella liriodendri*, which was also annotated using Helixer was found to have 16,580 protein-coding genes (Mongue et al. 2024), which is comparable to the number found in *N. viridis*. Further RNA evidence will allow us to confirm our estimated gene predictions for *N. viridis*.

### Repeat Masking

We found that repetitive elements accounted for 91,843,934 bases, or 32.40% of the sequence. A more in-depth breakdown of repeat content can be found in **Table 2**. In comparison to other species that have been sequenced to a similar level, *N. viridis* has a comparable repeat content to the cotton mealybug, *Phenacoccus solenopsis*, which has a ∼290Mb genome and ∼38% repetitive sequence (Li et al. 2020) In contrast, the larger soft scale *Toumeyella liriodendri* has a roughly 530Mb genome with 64.69% of the sequence being repetitive (Mongue et al. 2024), nearly double that of *N. viridis*, and consistent with a pattern of larger genomes having more repetitive elements (Yuan et al. 2024).

**Table 1.**
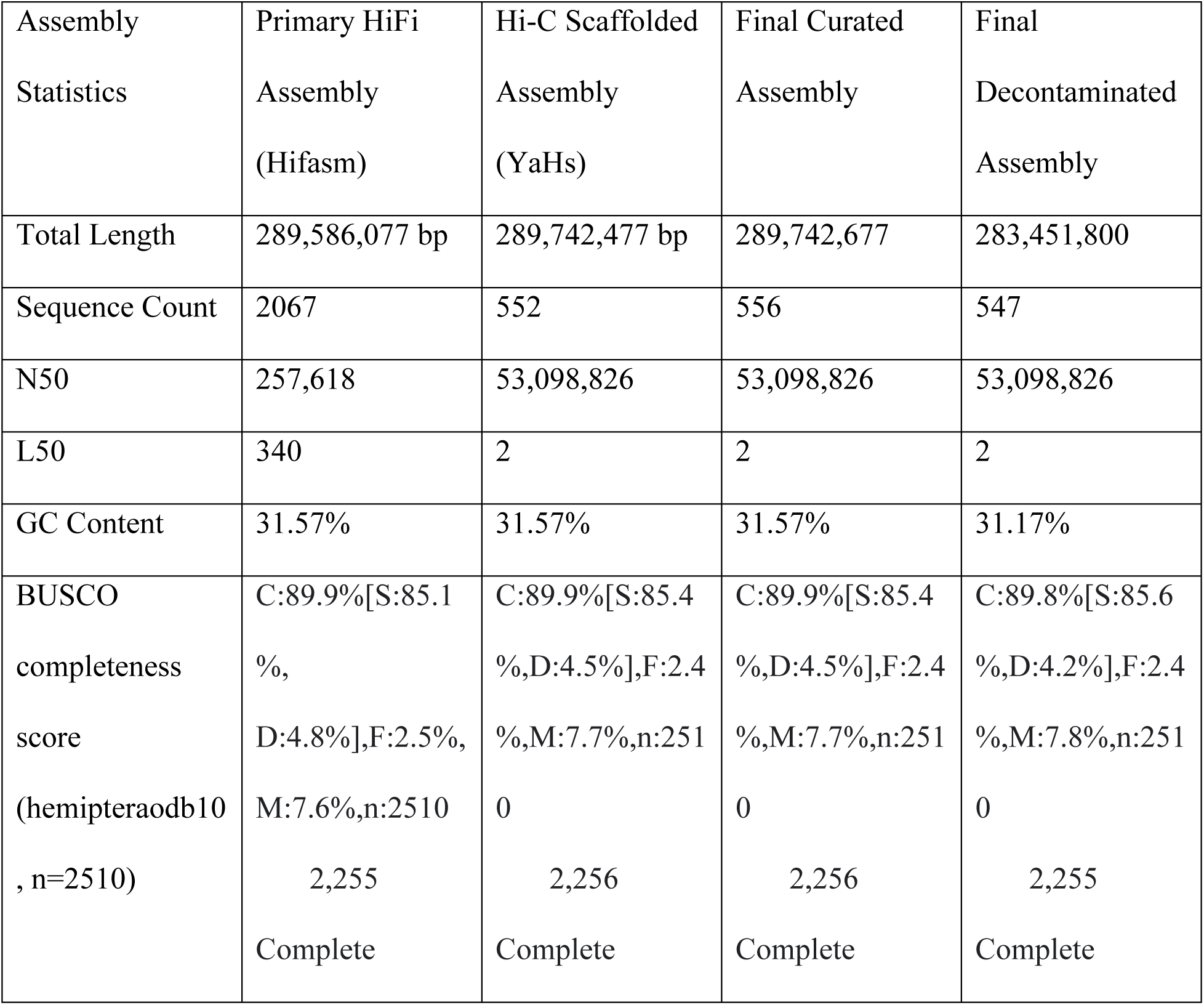
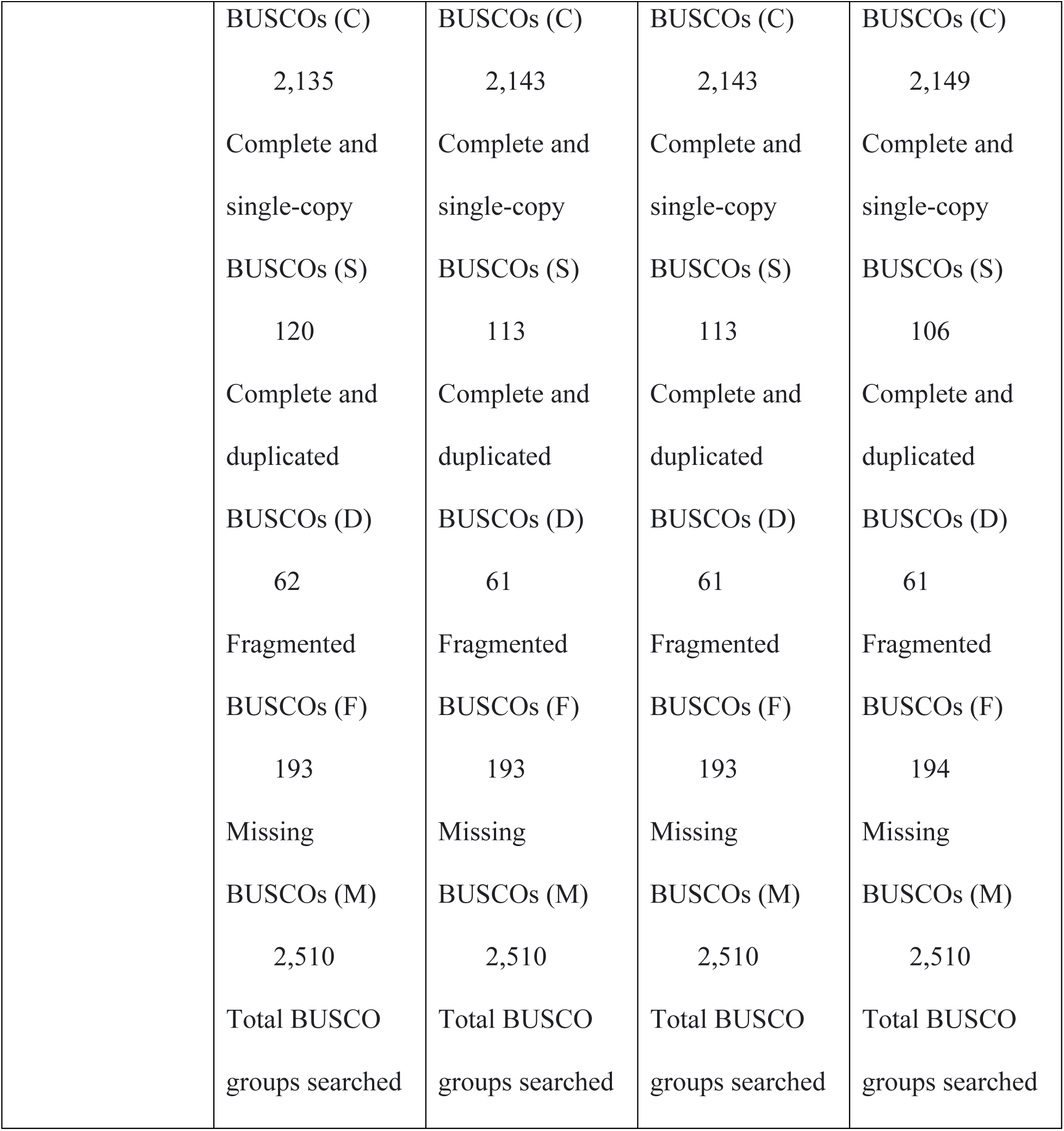
Assembly statistics of our primary HiFi assembly, Hi-C scaffolded assembly, final curated assembly, and the final decontaminated assembly. Details include assembly size, contiguity, and BUSCO completeness.

| Assembly Statistics | Primary HiFi Assembly (Hifasm) | Hi-C Scaffolded Assembly (YaHs) | Final Curated Assembly | Final Decontaminated Assembly |
| --- | --- | --- | --- | --- |
| Total Length | 289,586,077 bp | 289,742,477 bp | 289,742,677 | 283,451,800 |
| Sequence Count | 2067 | 552 | 556 | 547 |
| N50 | 257,618 | 53,098,826 | 53,098,826 | 53,098,826 |
| L50 | 340 | 2 | 2 | 2 |
| GC Content | 31.57% | 31.57% | 31.57% | 31.17% |
| BUSCO completeness score (hemipteraodb10, n=2510) | C:89.9%[S:85.1%,<br>D:4.8%],F:2.5%,<br>M:7.6%,n:2510<br>2,255<br>Complete | C:89.9%[S:85.4%,<br>D:4.5%],F:2.4%,<br>M:7.7%,n:2510<br>2,256<br>Complete | C:89.9%[S:85.4%,<br>D:4.5%],F:2.4%,<br>M:7.7%,n:2510<br>2,256<br>Complete | C:89.8%[S:85.6%,<br>D:4.2%],F:2.4%,<br>M:7.8%,n:2510<br>2,255<br>Complete |

|  |  |  |  |
| --- | --- | --- | --- |
| BUSCOs (C) | BUSCOs (C) | BUSCOs (C) | BUSCOs (C) |
| 2,135 | 2,143 | 2,143 | 2,149 |
| Complete and<br>single-copy | Complete and<br>single-copy | Complete and<br>single-copy | Complete and<br>single-copy |
| BUSCOs (S) | BUSCOs (S) | BUSCOs (S) | BUSCOs (S) |
| 120 | 113 | 113 | 106 |
| Complete and<br>duplicated | Complete and<br>duplicated | Complete and<br>duplicated | Complete and<br>duplicated |
| BUSCOs (D) | BUSCOs (D) | BUSCOs (D) | BUSCOs (D) |
| 62 | 61 | 61 | 61 |
| Fragmented | Fragmented | Fragmented | Fragmented |
| BUSCOs (F) | BUSCOs (F) | BUSCOs (F) | BUSCOs (F) |
| 193 | 193 | 193 | 194 |
| Missing | Missing | Missing | Missing |
| BUSCOs (M) | BUSCOs (M) | BUSCOs (M) | BUSCOs (M) |
| 2,510 | 2,510 | 2,510 | 2,510 |
| Total BUSCO<br>groups searched | Total BUSCO<br>groups searched | Total BUSCO<br>groups searched | Total BUSCO<br>groups searched |

**Table 2.** Summary of the masked repeats in the *N. viridis* genome.

| Repetitive element | Count | Sequence length (bp) | Percentage of sequence |
| --- | --- | --- | --- |
| Retroelements | 19,367 | 8,797,510 | 3.10 |
| SINEs: | 11 | 729 | 0.00 |
| LINEs: | 8,659 | 3,578,092 | 1.26 |
| L2/CR1/Rex | 194 | 74187 | 0.03 |
| R1/LOA/Jockey | 6,426 | 3,007,738 | 1.06 |
| R2/R4/NeSL | 106 | 13,176 | 0.00 |
| RTE/Bov-B | 407 | 139,248 | 0.05 |
| L1/CIN4 | 3 | 210 | 0.00 |
| LTR elements: | 10,697 | 5,218,689 | 1.84 |
| BEL/Pao | 1,080 | 553,216 | 0.20 |
| Ty1/Copia | 2,605 | 1,592,414 | 0.56 |
| Gypsy/DIRS1 | 2,526 | 2,053,227 | 0.72 |
| Retroviral | 162 | 18,065 | 0.01 |
| DNA transposons | 13,454 | 3,709,106 | 1.31 |
| hobo-Activator | 1,560 | 311,013 | 0.11 |
| Tc1-IS630-Pogo | 2,104 | 472,297 | 0.17 |
| MULE-MuDR | 465 | 112,103 | 0.04 |
| PiggyBac | 561 | 171,750 | 0.06 |
| Tourist/Harbinger | 168 | 56,109 | 0.02 |
| Other (Mirage, P-element, Transib) | 95 | 48,375 | 0.02 |
| Rolling-circles | 8065 | 1,475,777 | 0.52 |
| Unclassified: | 314,860 | 79,337,318 | 27.99 |
| Total interspersed repeats: |  | 91,843,934 | 32.40 |
| Satellites: | 17 | 1,159 | 0.00 |
| Simple repeats: | 136,474 | 4,512,679 | 1.59 |
| Low complexity: | 29,952 | 1,423,408 | 0.50 |

### Endosymbionts

In the blobplot visualization of our final curated assembly seen in Figure 3, we identified 5 chromosomal scaffolds as seen in the large, dark maroon color classified as “Arthropoda”. We also saw scaffolds of potential bacterial sequences, colored in teal, and labeled as “Pseudomonadota”. These two groups differ not only in coverage but the GC proportion as well, with the bacterial segments having a higher coverage and a higher GC proportion, around 0.55 compared to the chromosomal scaffolds at a proportion around 0.31. Higher coverage suggests that more of these molecules were sequenced than for the core mealybug genome. Some variation in sequence coverage is random and natural to the sequencing process, but the higher pattern suggests that these sequences were in higher copy number than the core genome, as expected of the numerous bacterial cells that exist within each mealybug cell in their symbiotic tissue, the bacteriome organ (Husnik and McCutcheon 2016). Thus, these differences are suggestive, but not conclusive of endosymbiont or other co-biont sequencing. To attempt to confirm the status and identity of endosymbionts, we further explored sequences as follows.

**Fig. 3.**
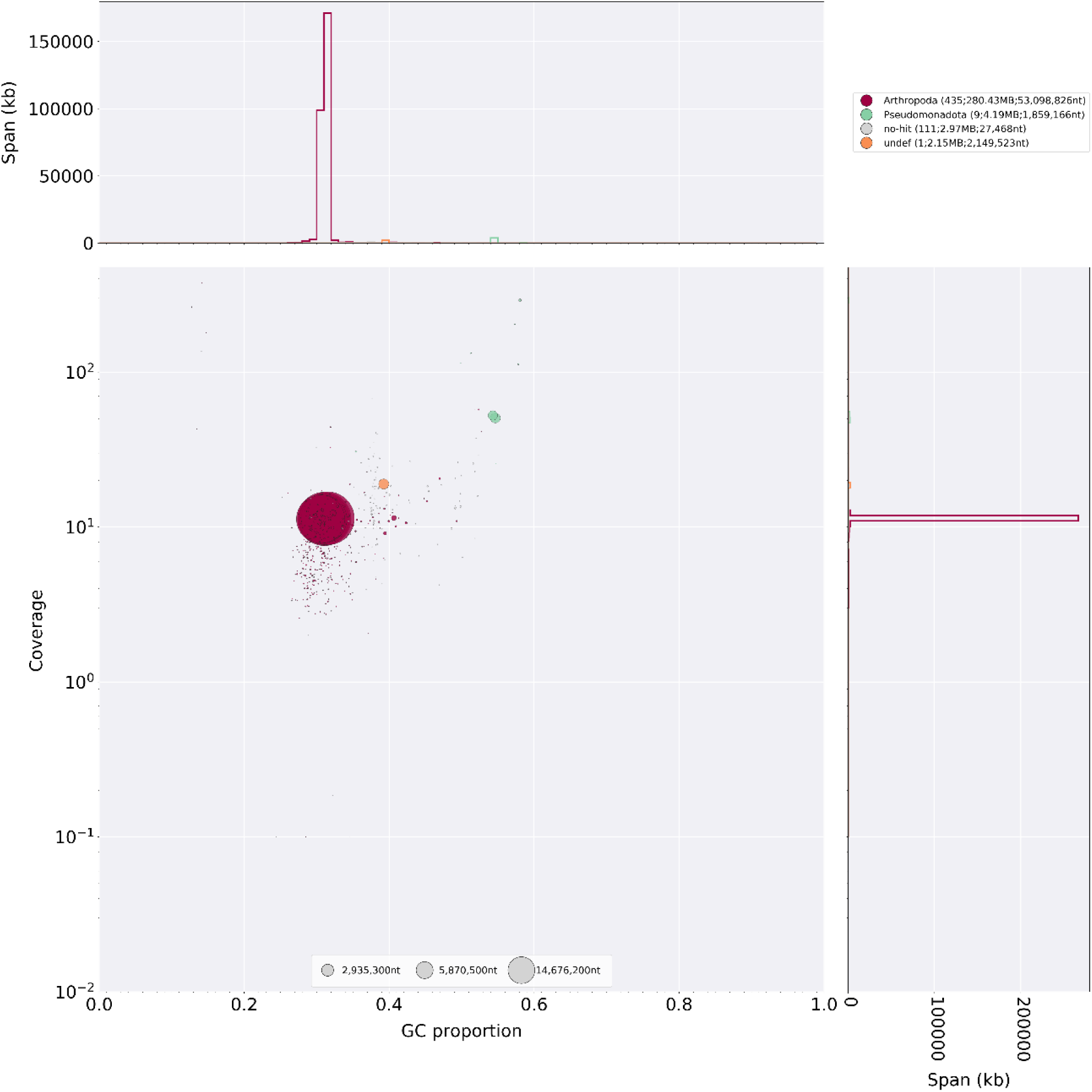
Blobplot visualization of the final curated assembly of *Nipaecoccus viridis*. Scaffolds are identified by the classification level of “Orders” by color and further can be examined for their differences in coverage (y-axis) and GC content (x-axis). Blob sizes are proportional to scaffold length. Endosymbiont sequences are expected to differ from the core mealybug genome in coverage, GC content, and taxonomic identification.

In the primary genome assembly, we identified a total of 13 potential bacterial sequences, with 3 being assembled into circular contigs (**Table 3**, left), as expected of most bacterial chromosomes. In manual BLAST searching of NCBI’s database, these hit to *Tremblaya,* and 2 different *Sodalis* species. Specifically, *Tremblaya princeps* is a known endosymbiont of other members of Pseudococcidae, including *Plannococcus citri* and *Pseudococcus longispinus* (Husnik and McCutcheon 2016). *Sodalis* species have also been known to be endosymbionts of mealybug species, but have also been detected within other arthropods such as tsetse flies; more specifically it seems that *Sodalis glossinidus* are more routinely and frequently acquired, which then is passed from one individual to another (Gatehouse et al. 2012). In tracking these contigs into our scaffolded assembly, all three sequences were recovered in the curated assembly as separate scaffolds (i.e., not assembled with other sequences). With this evidence taken together, these scaffolds represent the highest-confidence endosymbiont sequences.

**Table 3.** Identification of the endosymbiotic bacteria found within the genome assembly of *N. viridis.* Additional information includes the contig location within the primary assembly, scaffold location of the contig, and length of the bacterial segment. Contig names end with either “c” for circular or “l” for linear based on primary assembly.

| BLAST<br>Identification | BLAST<br>Query | Percent<br>Identity | Primary<br>assembly | Scaffold | Length<br>(bp) | GC<br>Content | Coverage<br>(x) | Final<br>Assignment |
| --- | --- | --- | --- | --- | --- | --- | --- | --- |

|  | Coverage |  | contig |  |  | (%) |  |  |
| --- | --- | --- | --- | --- | --- | --- | --- | --- |
| *Sodalis praecaptivus* | 87% | 97.84% | ptg00004<br>2c | scaffold<br>_8 | 1,859,<br>166 | 54.3 | 53 | Putative symbiont, removed from assembly |
| *Sodalis glossinidius* | 59% | 92.58% | ptg00015<br>2c | scaffold<br>_7 | 2,030,<br>015 | 54.7 | 51 | Putative symbiont, removed from assembly |
| *Candidatus Tremblaya princeps* | 100% | 5.64% | ptg00025<br>9l | scaffold<br>_25 | 130,20<br>3 | 58.0 | 291 | Putative symbiont, removed from assembly |
| *Proteus appendicitidis* | 5% | 73.65% | ptg00029<br>6l | scaffold<br>_6 | 2,149,<br>523 | 39.2 | 19 | Putative symbiont, removed from assembly |
| *Planococcus citri* | 17% | 92.21% | ptg00031<br>6l | scaffold<br>_3 | 110,26<br>2 | 31.0 | 11 | Likely<br>horizontal<br>gene transfer,<br>retained |
| *Planococcus citri* | 2% | 88.68% | ptg00079<br>8l | scaffold<br>_2 | 86,036 | 31.5 | 11 | Likely<br>horizontal<br>gene transfer,<br>retained |
| *Burkholderia pseudomallei* | 0% | 89.61% | ptg00108<br>8l | scaffold<br>_73 | 53,456 | 49.3 | 11 | Unclear<br>assignment,<br>retained |
| *Candidatus Tremblaya princeps* | 90% | 94.90% | ptg00114<br>5l | scaffold<br>_135 | 35,172 | 57.8 | 113 | Putative<br>symbiont,<br>removed from assembly |
| *Candidatus Tremblaya princeps* | 99% | 95.74% | ptg001360c | scaffold_268 | 20,985 | 57.3 | 204 | Putative symbiont, removed from assembly |
| *Sodalis glossinidius* | 34% | 92.58% | ptg001498l | scaffold_162 | 30,728 | 51.3 | 133 | Putative symbiont, removed from assembly |
| *Candidatus Sodalis pierantonius* | 31% | 93.59% | ptg001522l | scaffold_270 | 20,905 | 49.8 | 114 | Putative symbiont, removed from assembly |
| *Arsenophonus nasoniae* | 6% | 74.75% | ptg001825l | scaffold_83 | 47,879 | 35.4 | 30 | Unclear assignment, retained |
| *Sodalis*<br>*glossinidius* | 66% | 93.05% | ptg00199<br>51 | scaffold<br>_464 | 14,180 | 54.7 | 26 | Putative<br>symbiont,<br>removed from<br>assembly |

A further five contigs had BLAST hits to symbionts, did not assemble further with Hi-C data, but were initially assembled as linear sequences. Although we expect circular bacterial assemblies, the weight of evidence suggests that these are (partial) bacterial sequences as well.

We excluded these eight scaffolds from the core genome for archiving purposes as they are almost certainly not *N. viridis* sequences. We include them as a supplemental fasta file for those interested but do not consider the endosymbionts in greater detail here.

Of the remaining five potential bacterial contigs, we found two that assembled into the chromosomal scaffolds (scaffolds 1 – 5). The contig “ptg000798l” was placed on scaffold_2 and “ptg000316l” was found in scaffold_3 based on the Hi-C contact with other core genomic sequences. When BLASTing these sequences against NCBI’s database, the top hits were not bacterial species, but rather another mealybug species, *Planococcus citri*. The similarity to this mealybug and contact with *N. viridis* sequences suggests that these sequences are part of the core mealybug genome. Thus, the similarity to bacteria in the Blobplot could represent horizontal transfer of genes from bacteria to insect, which has been reported in other mealybugs (Husnik and McCutcheon 2016). As such, we left these sequences intact as part of the mealybug genome.

Finally, two putative contigs, “ptg001088l” and “ptg001825l” were found to be identified as the bacterial species, *Burkholderia pseudomallei* and *Arsenophonus nasoniae* respectively. However, in the manual BLAST step, both contig sequences were found to have very low (<10%) query coverages in the identification of these species and both had passed NCBI’s contaminant screening on genome upload, so we did not remove these sequences from our core genome assembly.

## Conclusions

Our report is a chromosome-level assembly for *Nipaecoccus viridis*, an important widespread invasive agricultural pest. This assembly produces the necessary resources for further research of the applied side to expand our knowledge about its path of invasive spread, not just in Florida, but to provide a means of comparison to other worldwide records of this pest. Furthermore, this genome provides the baseline steps to other key features within this insect such as a deeper analysis of their unique sex determination system known as Paternal Genome Elimination (PGE), and a closer look at the unique composition of endosymbiotic bacteria housed within this insect.

## Data Availability

Data will be publicly available through the National Library of Medicine: National Center for Biotechnology Information (NCBI) and the accession numbers for the genome assembly and raw sequence data can be found in **Table 4**. The annotation is hosted at https://github.com/TLiesenfelt/NiviGenome, and potential symbiont and contaminant sequences are provided as a supplementary file. Reviewer link to genome assembly data: https://dataview.ncbi.nlm.nih.gov/object/PRJNA1104600?reviewer=mh36o29uni5j5sq7bvatuive09

**Table 4:** Location of assembly data for *N. viridis*.

| Data | Accession (Bioproject: PRJNA1104600) |
| --- | --- |
| PacBio HiFi Reads | SRR28800798 |
| Illumina Hi-C Reads | SRR28800797 |
| Assembly | SUB14816508 |

